# The spread of chloramphenicol-resistant *Neisseria meningitidis* in Southeast Asia

**DOI:** 10.1101/2019.12.13.872499

**Authors:** Elizabeth M. Batty, Tomas-Paul Cusack, Janjira Thaipadungpanit, Wanitda Watthanaworawit, Verena Carrara, Somsavanh Sihalath, Jill Hopkins, Sona Soeng, Clare Ling, Paul Turner, David A.B. Dance

## Abstract

Invasive disease caused by *Neisseria meningitidis* is a significant health concern globally, but our knowledge of the prevailing serogroups, antimicrobial susceptibility patterns, and genetics of *N. meningitidis* in Southeast Asia is limited. Chloramphenicol resistance in *N. meningitidis* has rarely been reported, but was first described in isolates from Vietnam in 1998. Using whole-genome sequencing of meningococcal isolates from 18 patients collected between 2007 and 2018 from diagnostic microbiology laboratories in Cambodia, Thailand and the Lao People’s Democratic Republic (Laos), of which eight were non-susceptible to chloramphenicol, we report the spread of this chloramphenicol-resistant lineage of *N. meningitidis* across Southeast Asia. Strains resistant to penicillin, tetracycline, and ciprofloxacin were also observed, including a chloramphenicol-resistant strain from the previously-described lineage which has acquired penicillin and ciprofloxacin resistance, and most isolates were of serogroup B. This study suggests that chloramphenicol-resistant *N. meningitidis* is more widespread than previously thought.

## Introduction

Invasive disease caused by *Neisseria meningitidis* is an important global health concern, being associated with significant rates of mortality and long-term sequelae (Vyse et al. 2013). The burden of meningococcal disease in Southeast Asia is poorly described, with few data on prevailing serogroups and antimicrobial susceptibility patterns (Vyse et al. 2011). The Mahidol-Oxford Tropical Health Network (MORU) includes diagnostic microbiology laboratories in Thailand, Laos and Cambodia, where *N. meningitidis* is an infrequently isolated pathogen. Detection of a chloramphenicol-resistant strain of *N. meningitidis* at our laboratory site in Laos in November 2017 prompted a review of published reports of chloramphenicol resistance in *N. meningitidis* and review of chloramphenicol susceptibility amongst meningococcal isolates in our laboratory network. Although no longer standard therapy for meningococcal meningitis, chloramphenicol is commonly used in beta-lactam intolerant patients and is recommended as an alternative agent in international guidelines for treatment of meningococcal meningitis (Tunkel et al. 2004; McGill et al. 2016), and therefore resistance to this agent remains an issue of clinical importance.

Chloramphenicol resistance in *N. meningitidis* has rarely been reported, first being described in 1998 in 11 isolates from Vietnam and one isolate from France, due to the presence of a chloramphenicol transferase gene (*catP*) possibly derived from a *Clostridium perfringens* transposable element (Galimand et al. 1998). Two further isolates with the same *catP* insertion were later described in Australia (Shultz et al. 2003) amongst 1382 isolates. More recently, chloramphenicol resistance was reported in one of 2888 isolates in Brazil (Gorla et al. 2018), and a further highly resistant isolate was found in Vietnam in 2014 (Tran et al. 2019). Despite historical use of single-dose oily chloramphenicol for epidemic management of meningococcal meningitis (Fuller et al. 2003), we could find no reports of chloramphenicol resistance from Africa, although susceptibility information on African isolates is limited (Hedberg et al. 2009).

We identified chloramphenicol non-susceptibility in eight of 29 isolates of *N. meningitidis* in our network laboratories over an 11 year time period. Given the seemingly high frequency of chloramphenicol non-susceptibility amongst our *N. meningitidis* isolates, we aimed to characterise the genetic basis of this using whole genome sequencing. In addition, due to the scarcity of meningococcal data from Southeast Asia, we report serogroup data and phenotypic and genotypic susceptibilities to other routinely tested antimicrobial agents in 18 of these isolates that were available for sequencing.

## Methods

### Study sites

Clinical isolates from three laboratories were included. The Lao-Oxford-Mahosot Hospital-Wellcome Trust Research Unit (LOMWRU), Microbiology Laboratory, Mahosot Hospital, Vientiane, Laos provides diagnostic microbiology services to Mahosot Hospital as well as other hospitals in Vientiane and other provincial sites. The Microbiology Laboratory of the Shoklo Malaria Research Unit (SMRU), Mae Sot, Thailand supports healthcare provision to marginalised populations on the Thailand-Myanmar border. The Cambodia-Oxford Medical Research Unit (COMRU) provides diagnostic microbiology services to the Angkor Hospital for Children, a paediatric teaching hospital in Siem Reap, Cambodia. All laboratories routinely store clinically significant bacterial isolates at −80°C.

### Sample selection, organism identification and phenotypic antimicrobial susceptibility testing

Laboratory records were reviewed for all clinical isolates of *N. meningitidis* up until 31^st^ March 2018, with data available from year 2000 in LOMWRU, 2007 in SMRU and from 2013 in COMRU. A total of thirty-four isolates were identified (9 from LOMWRU, 18 from SMRU, 7 from COMRU), of which twenty-five were stored and viable. All had susceptibility data for chloramphenicol, penicillin, ciprofloxacin and ceftriaxone. *N. meningitidis* isolates were originally identified by API NH (bioMérieux, France). Antimicrobial susceptibility testing was performed by disk diffusion for chloramphenicol, ciprofloxacin and ceftriaxone, and Etest (bioMérieux, France) for penicillin, according to Clinical and Laboratory Standards Institute (CLSI) standards and breakpoints. All susceptibility testing was performed in duplicate. Before genotypic characterisation, isolates were subcultured on to chocolate agar from storage at −80°C and underwent confirmation of identity by MALDI-TOF (Vitek MS, bioMérieux) and susceptibility testing repeated and interpreted using the CLSI guideline from 2018 (Wayne, PA, Clinical and Laboratory Standards Institute 2018). Two isolates failed to confirm as *N. meningitidis*, and five isolates were removed as duplicate samples where a different tissue had also been sampled from the same patient, leaving 18 isolates for analysis.

### Patient-related metadata

Demographic data, clinical syndrome, antibiotic treatment and status on discharge were obtained from laboratory records and patient charts where available.

### Genotypic characterization

Sequencing was performed at the MORU laboratories in Bangkok, Thailand. Libraries were made from extracted DNA using the Illumina Nextera XT library preparation kit (Illumina, San Diego, CA, USA) and quantified on an Agilent BioAnalyser (Agilent Technologies, Santa Clara, CA, USA). The pooled libraries were sequenced on one run of an Illumina MiSeq sequencer using the v2 reagent kit to give 250bp paired-end reads. The genome assembly of chloramphenicol-resistant strain DuyDNT from Vietnam was obtained from Genbank under accession RPSF00000000.

Spades v3.11.1 (Bankevich et al. 2012) was used to assemble genomes from whole-genome sequencing. Abricate v0.8.10 (Seemann) was used to look for acquired resistance genes using the bundled ResFinder database (updated 2018-10-02) (Zankari et al. 2012). BLAST was used to search for specific antibiotic resistance genes. Prokka 1.13.3 was used to annotate the assembled genomes.

In-silico serotyping of the isolates was performed by retrieving the sequences of the genes in the capsule synthesis region of the capsule that are unique to each serotype (*csb, csc, csw* and *csy*) for the B, C, W and Y serogroups respectively, and determining their presence or absence using BLAST to search the genome assemblies.

Snippy v4.3.6 (Seemann) was used to call SNPs against the MC58 reference genome from raw sequence data and from the assembled DuyDNT genome. Initial phylogenetic trees were generated from the core SNP fasta files using IQ-TREE (Nguyen et al. 2015) under the GTR+F+ASC+R3model which was the best fit using BIC implemented in ModelFinder (Kalyaanamoorthy et al. 2017). The initial tree was used as input to Gubbins v2.3.1 (Croucher et al. 2015) to generate a phylogeny accounting for recombination.

## Results

### Phenotypic characteristics and patient-related data

Eighteen isolates of *N. meningitidis* (11 from Thailand, 5 from Cambodia, 2 from Laos) from 18 patients were included in the analysis (Table 1). Eight of 18 (44.4%) meningococcal isolates were non-susceptible to chloramphenicol (3 intermediate, 5 resistant). Four of the non-susceptible isolates were collected from Thailand, and two each from Cambodia and Laos. All isolates were susceptible to ceftriaxone, but 10/18 (55.5%) isolates had reduced susceptibility to penicillin, including one isolate which was resistant according to CLSI criteria (Table 1). Phenotypic ciprofloxacin resistance was detected in one isolate.

**Table 1.** Clinical and genetic information on the samples. SMRU, Shoklo Malaria Research Unit, Mae Sot, Thailand; LOMWRU, Lao-Oxford-Mahosot Hospital Wellcome Trust Research Unit, Vientiane, Laos; COMRU, Cambodia Oxford Medical Research Unit, Siem Reap, Cambodia CSF, cerebrospinal fluid; BC, blood culture CHL, chloramphenicol; CIP, ciprofloxacin; CRO, ceftriaxone; PEN, penicillin; TET, tetracycline S, susceptible; I, intermediate; R, resistant; MIC, minimum inhibitory concentration

| Sample ID | Unit | Specimen Date | Sample type | Age (years) | Sex | Phenotypic antibiotic susceptibility |  |  |  | Genotypic antibiotic resistance | Serogroup | MLST | Diagnosis |
| --- | --- | --- | --- | --- | --- | --- | --- | --- | --- | --- | --- | --- | --- |
|  |  |  |  |  |  | CHL | CIP | CRO | PEN (MIC) |  |  |  |  |
| NM01 | SMRU | 25/7/07 | CSF | 7 | M | I | S | S | I (0.12) | CHL, PEN | B | 1487 | Meningitis |
| NM03 | SMRU | 25/11/09 | BC | 1 | F | S | S | S | I (0.25) | PEN | B | 14488 | Sepsis |
| NM04 | SMRU | 15/11/10 | BC | <1 | M | S | S | S | S (0.06) |  | B | 14489 | Meningitis |
| NM06 | SMRU | 19/6/11 | BC | 0.2 | M | S | S | S | S (0.06) |  | B | 32 | Meningitis |
| NM07 | SMRU | 4/10/11 | BC | 0.5 | M | S | S | S | I (0.25) | PEN, CIP, TET | C | 3256 | Meningitis |
| NM09 | SMRU | 7/1/12 | BC | 0.2 | M | S | S | S | S (0.06) |  | B | 5604 | Meningitis |
| NM11 | SMRU | 18/1/12 | BC | 3 | M | I | S | S | S (0.06) | CHL | B | 14496 | Meningitis |
| NM12 | SMRU | 28/12/12 | BC | 0.3 | F | I | S | S | R (0.5) | CHL, PEN | B | 1576 | Sepsis |
| NM13 | SMRU | 13/5/13 | BC | 0.6 | M | R | S | S | S (0.06) | CHL | B | 1576 | Sepsis |
| NM14 | SMRU | 15/2/14 | BC | 0.3 | F | S | S | S | I (0.25) | PEN | B | 1145 | Sepsis |
| NM15 | SMRU | 19/12/14 | BC | 0.3 | M | S | S | S | I (0.25) | PEN | B | 41 | Sepsis |
| NM16 | LOMWRU | 10/11/17 | CSF | 60 | F | R | S | S | I (0.12) | CHL | B | 1576 | Meningitis |
| NM18 | COMRU | 13/10/13 | BC | 0.14 | F | R | R | S | I (0.12) | CHL, CIP, PEN | B | 1576 | Meningitis/ Sepsis |
| NM19 | COMRU | 25/4/14 | BC | 1.84 | F | S | S | S | S (0.03) |  | C | 14503 | Sepsis |
| NM20 | COMRU | 31/3/15 | BC | 0.91 | F | R | S | S | S (0.06) | CHL, PEN | B | 11005 | Pneumonia/ Sepsis |
| NM21 | COMRU | 24/7/16 | BC | 0.07 | M | S | S | S | S (0.016) |  | B | 12811 | Meningitis/ Sepsis |
| NM23 | COMRU | 22/8/17 | BC | 0.33 | F | S | S | S | I (0.12) | PEN | B | 14507 | Meningitis |
| NM25 | LOMWRU | 21/3/18 | BC | 0.25 | M | R | S | S | I (0.25) | CHL, PEN | B | 1576 | Meningitis |

Despite COMRU being the only site with a purely paediatric population, 17/18 patients were children under 8 years old, with a median age of approximately 4 months (one patient <1 year old with no recorded date of birth was attributed the age of 1 year for the purposes of the calculation). Most patients presented with meningitis and/or sepsis, all patients with known antibiotic treatment (14/18) were treated with ceftriaxone, and all of the 16 patients with known outcome were discharged alive following clinical improvement.

### Genotypic characterization

All 18 isolates underwent sequencing. 0.4-2.4 million sequence reads were obtained from each isolate, giving 42-233X coverage of the genome. The genomes were assembled in 133-248 contigs, with a total length of 2,124,885-2,217,307bp and n50 between 40.4-82.5kb.

We determined the serotypes of the isolates from the genome assemblies by looking for the presence of the unique capsule synthesis genes for the B, C, W and Y serogroups which commonly cause invasive disease (Harrison et al. 2013). Two isolates showed the presence of the *csc* gene, indicating they are serogroup C, while the other sixteen isolates, including all the resistant isolates, had the *csb* gene, indicating they are serogroup B. No matches for the serogroup W or Y genes could be found.

We determined the sequence types of the isolates in our sample set (Table 1). Of the chloramphenicol non-susceptible isolates, five were ST1576, one was ST11005, and two isolates had novel sequence types (assigned as ST14488 and ST14507). ST1576 and ST11005 differ only at the *gdh* locus, and the two novel sequence types also differ from ST1576 at only one locus. None of the chloramphenicol susceptible isolates were from ST1576 or ST11005, but they were from multiple sequence types, including three isolates with novel sequence types (assigned as ST14496, ST14503, and ST14507). Two isolates from Thailand were ST41 and ST1145, and one was from ST32. These sequence types are part of the ST41/44 and ST32 hypervirulent clonal complexes commonly seen in invasive disease (Racloz and Luiz 2010). The other isolates could not be assigned to a clonal complex.

While searching for the presence of antimicrobial resistance genes in the isolates, we identified a 623bp *catP* chloramphenicol transferase gene with the same flanking sequences in all eight chloramphenicol resistant isolates. It had previously been shown that a *catP* gene was present in the highly chloramphenicol-resistant DuyDNT isolated in Vietnam in 2014, and the same *catP* gene and flanking regions also present in this isolate. By alignment of the region surrounding the *catP* gene to the MC58 reference genome, it could be determined that the chloramphenicol resistance gene was found inserted between genes NMB1351 and NMB1352. The *catP* gene is identical to the *catP* gene previously reported in France (Galimand et al. 1998) and a comparison of the sequences surrounding the insertion show that it is inserted in the same position in the genome (Supplementary Figure 1). The insertion is 986bp identical to the Tn4451 transposon from *Clostridium perfringens*, but contains only the *catP* gene and no complete flanking genes.

Ten of our isolates had reduced susceptibility or resistance to penicillin, including two of the chloramphenicol-resistant isolates (NM18 and NM25). We extracted the *penA* gene sequence to look for the five amino acid changes known to confer reduced susceptibility to penicillin (Taha et al. 2007). Ten isolates had all five amino acid changes, while the other twelve isolates had none of the five changes. Concordance between the phenotypic and genotypic susceptibility was not complete – one of the isolates with reduced susceptibility phenotypically did not display the amino acid changes, while one isolate with the *penA* changes did not show phenotypic reduced susceptibility. The isolate determined to be resistant to penicillin had the same *penA* allele as an isolate with only reduced susceptibility. The bla_ROB-1_ gene is also known to confer penicillin resistance in *N. meningitidis* (Tsang et al. 2019), but this gene was not present in any of our strains.

Isolate NM07 from Thailand was found to contain the tetracycline resistance gene *tetB*. NM07 and NM18, carry the *gyrA* Thr91 → Ile mutation known to cause ciprofloxacin resistance (Enríquez et al. 2008; Skoczynska et al. 2008), although only NM18 was phenotypically resistant to ciprofloxacin.

We generated a phylogeny of the 18 genomes from this study and the previously-reported DuyDNT chloramphenicol-resistant isolate (Figure 1) after removing recombinant SNPs, as *N. meningitidis* is highly recombinogenic (Feil et al. 1999). This shows that the chloramphenicol-resistant isolates from all four countries are in a single clade, with the isolates from each country clustered together within this clade. Chloramphenicol-susceptible isolates from Thailand and Cambodia are not separated phylogenetically, and the serogroup C isolates (NM07 and NM19) are not clustered separately from the serogroup B isolates.

**Figure 1.**
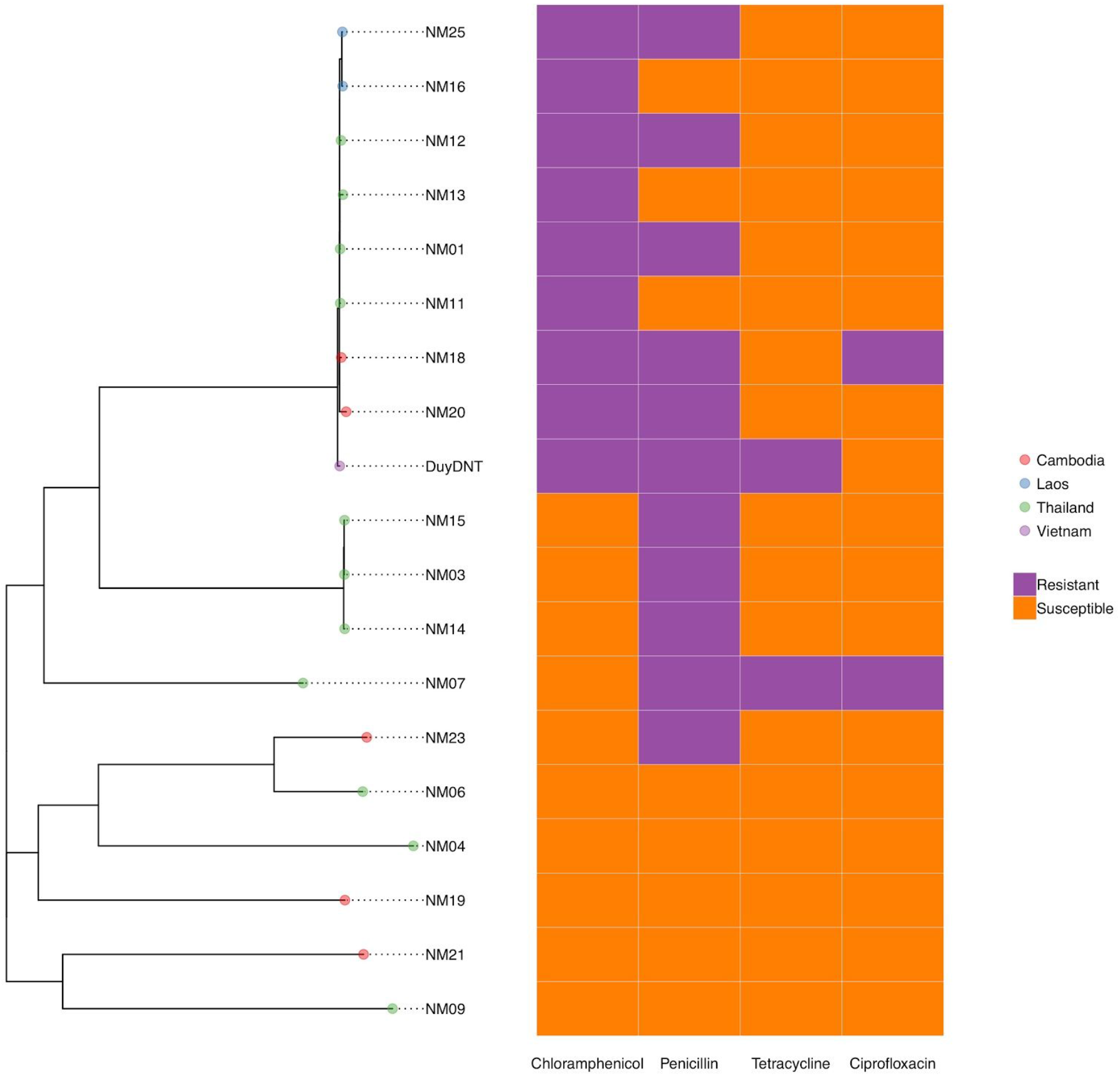
A phylogeny of the 19 *Neisseria meningitidis* strains showing country of origin and resistance genotypes.

## Discussion

We describe phenotypic and genotypic characteristics of 18 isolates of *N. meningitidis* from three countries in Southeast Asia isolated over an 11 year period. Most strikingly, chloramphenicol non-susceptibility, although rarely reported in the literature, was detected in a significantly higher proportion (44.4%) of isolates compared to studies from other regions reporting on susceptibilities of large numbers of isolates, suggesting this is proportionately more common in Southeast Asia. The identical *catP* insertion across the isolates in this study and the previously reported isolates from Vietnam and Australia suggest that chloramphenicol resistance in *N. meningitidis* was acquired once in the ST1576 lineage, and that lineage has now spread across Southeast Asia. The wide range of MICs determined across isolates with the same identified chloramphenicol resistance gene suggest that there may be further determinants of resistance in these genomes which have yet to be identified.

A high proportion of our isolates (55.5%) displayed reduced susceptibility to penicillin, 9/10 of which had genetic markers known to confer reduced penicillin susceptibility. However, a penicillin-susceptible isolate also carried these genetic markers, although the MIC in this isolate was close to the intermediate breakpoint. Similarly, the one isolate showing phenotypic ciprofloxacin resistance had a known mutation conferring resistance to ciprofloxacin, but another isolate with the same mutation did not display phenotypic resistance. There may also be further genetic determinants of resistance in these strains, as we have limited knowledge of *N. meningitidis* genomes in Southeast Asia.

We also identified isolates resistant to multiple antibiotics including an isolate resistant to chloramphenicol and ciprofloxacin with reduced susceptibility to penicillin, and the DuyDNT isolate is susceptible to ciprofloxacin but carries a tetracycline resistance gene. The first reported chloramphenicol resistant *N. meningitidis* isolates were susceptible to penicillin, tetracycline, and quinolones (Galimand et al. 1998), suggesting that this lineage has acquired resistance to multiple antibiotics during its spread.

Our isolates were predominantly serogroup B, with only two patients with strains from serogroup C. This is consistent with previous reports of serogroup B isolates causing sporadic disease in Thailand (Pancharoen et al. 2000; Borrow et al. 2017) while very little information is available about common serogroups in Laos and Cambodia. The serogroup B and serogroup C strains were not separated in the phylogeny, suggesting a capsule switch may have occurred in these strains.

While some of the isolates we identified come from hypervirulent clonal complexes commonly implicated in invasive disease, the majority of them could not be assigned to one of these lineages, and many of these isolates were from novel sequence types. This suggests predominantly sporadic infection arising from the nasopharyngeal flora rather than an ongoing serogroup B epidemic.

There are several limitations of this study. The study involved only three sites and a relatively small number of isolates, reflecting the relative rarity of invasive meningococcal disease in our populations, giving only a snapshot of the epidemiology across the region. Furthermore, only culture-positive isolates were included, which may have underestimated the true number of cases and selected for more resistant isolates given the seemingly high prevalence of pre-hospital antibiotic treatment regionally (Khennavong et al. 2011), although relatively few additional cases were identified by routine CSF PCR for *N. meningitidis* in our laboratories. Nevertheless, our results suggest that resistance to chloramphenicol is relatively common in meningococcal isolates from Southeast Asia. This remains clinically important given the role of chloramphenicol as a common second-line agent for treatment of meningococcal meningitis. *N. meningitidis* which is non-susceptible to third-generation cephalosporins (Deghmane et al. 2017) has been recently reported in France, with a *penA* allele which may have been acquired from *Neisseria gonorrhoeae*, raising the possibility that recombination with extensively drug-resistant *N. gonorrhoeae* could result in the spread of further drug resistance in *N. meningitidis*. Our phenotypic and genotypic characterisation of isolates from three countries in Southeast Asia will add to the limited published data of the epidemiology of meningococcal disease in Southeast Asia, and aid future surveillance of drug-resistant isolates.

## Acknowledgements

This publication made use of the PubMLST website (https://pubmlst.org/) developed by Keith Jolley (Jolley and Maiden 2010) and sited at the University of Oxford. The development of that website was funded by the Wellcome Trust. We acknowledge the laboratory staff who assisted with this project, including Jiraporn Tangmanakit and Pattaraporn Hinfonthong at SMRU. We also thank Dr Manivanh Vongsouvath, Mahosot Microbiology Laboratory Director, Professor Paul Newton, Director of LOMWRU, and the staff of Mahosot Hospital and Lao Ministry of Health for their help and support. This work was funded by a Wellcome Trust grant (106698) to the Thailand Major Overseas Programme. The funders had no role in study design, in the collection, analysis and interpretation of data; in the writing of the manuscript; and in the decision to submit the manuscript for publication.

## Author Contributions

DD and PT conceived the project. EMB and TPC performed data analysis and drafted the manuscript. JT, WW, VC, SS, JH, and CL performed laboratory experiments. EMB, TPC, JH, CL, PT and DD edited the final manuscript. All authors reviewed and approved the final manuscript.

## Data Availability

Raw sequence data has been deposited in the European Nucleotide Archive as project PRJEB30968. Novel sequence types and isolates have been deposited in PubMLST.

