## Supplementary Figure 1 for "The spread of chloramphenicol-resistant *Neisseria meningitidis* in Southeast Asia"

```
a)
NM01      TGTATGTTTATTAAATTCTAAATCAATAATAATATTCCCAATCAGAGCTT
NM11      TGTATGTTTATTAAATTCTAAATCAATAATAATATTCCCAATCAGAGCTT
NM12      TGTATGTTTATTAAATTCTAAATCAATAATAATATTCCCAATCAGAGCTT
NM13      TGTATGTTTATTAAATTCTAAATCAATAATAATATTCCCAATCAGAGCTT
NM16      TGTATGTTTATTAAATTCTAAATCAATAATAATATTCCCAATCAGAGCTT
NM18      TGTATGTTTATTAAATTCTAAATCAATAATAATATTCCCAATCAGAGCTT
NM20      TGTATGTTTATTAAATTCTAAATCAATAATAATATTCCCAATCAGAGCTT
NM25      TGTATGTTTATTAAATTCTAAATCAATAATAATATTCCCAATCAGAGCTT
DuyDNT    TGTATGTTTATTAAATTCTAAATCAATAATAATATTCCCAATCAGAGCTT
MC58      TGTATGTTTATTAAATTCTAAATCAATAATAATATTCCCA----------

b)
NM01      GTTAGTACAAAGACCTTGTGTTTCTTTTTAACCAATATTTCATATATATC
NM11      GTTAGTACAAAGACCTTGTGTTTCTTTTTAACCAATATTTCATATATATC
NM12      GTTAGTACAAAGACCTTGTGTTTCTTTTTAACCAATATTTCATATATATC
NM13      GTTAGTACAAAGACCTTGTGTTTCTTTTTAACCAATATTTCATATATATC
NM16      GTTAGTACAAAGACCTTGTGTTTCTTTTTAACCAATATTTCATATATATC
NM18      GTTAGTACAAAGACCTTGTGTTTCTTTTTAACCAATATTTCATATATATC
NM20      GTTAGTACAAAGACCTTGTGTTTCTTTTTAACCAATATTTCATATATATC
NM25      GTTAGTACAAAGACCTTGTGTTTCTTTTTAACCAATATTTCATATATATC
DuyDNT    GTTAGTACAAAGACCTTGTGTTTCTTTTTAACCAATATTTCATATATATC
MC58      ---------------------------TTAACCAATATTTCATATATATC
```

Supplementary Figure 1. A sequence alignment of the nine chloramphenicol resistant and intermediate strains and the MC58 reference genome, showing the a) upstream and b) downstream flanking regions of the *catP* insertion which match the previously-reported insertion.
